# Hunger neurons track available food locations during foraging and spatial memory recall

**DOI:** 10.64898/2026.07.28.741066

**Authors:** Anna Gruzdeva, Jamien Shea, Daniel Shi, Antonio Fernandez-Ruiz, Azahara Oliva, Nilay Yapici

## Abstract

Foraging requires animals to integrate metabolic needs with knowledge of where resources are located. Agouti-related peptide (AgRP) neurons in the hypothalamus are central regulators of appetitive behaviors, yet their role in foraging remains poorly understood. Here, we used fiber photometry to record AgRP neuron activity in freely foraging mice and found that it dynamically tracks spatial proximity to food. AgRP neuron activity decreases progressively as mice approach the food source and increases as they move away, forming a gradient that scales with distance. Notably, this proximity signal emerges only in fasted mice once food is discovered, is specific to the accessible rather than to inaccessible sources, and persists where a source was previously available. Together, our findings reveal that AgRP neuron activity reflects distance to food in a dynamic, experience-dependent manner, extending what these neurons convey beyond internal need and food-related cues to include learned spatial information.

**Highlights:**

- AgRP neuron activity reflects spatial distance to food sources during foraging
- AgRP neuron activity gradually decreases on approach to food and increases during departure
- AgRP distance signal depends on metabolic state, and its recall requires visual cues
- AgRP neuron activity tracks food source availability and its location after source removal

## Introduction

Most animals forage in dynamic environments where food sources are patchy, and their availability is unpredictable^1,2^. Successful foraging therefore involves several neural processes: learning and remembering where food is located, navigating efficiently toward it^3^, and choosing resources that match the animal’s current metabolic need^4,5^. Hunger is a powerful driver for initializing these foraging processes: it shifts behavioral priorities toward food-seeking, heightens attention to food-predictive cues, and biases exploration toward locations previously associated with food availability. Decades of work have identified hypothalamic circuits as key regulators of food intake and metabolism, with Agouti-related peptide (AgRP)-expressing neurons in the arcuate nucleus (ARC) serving as a central node in this energy homeostasis system^6–14^. AgRP neurons are activated by food deprivation^9,15^, drive feeding even in satiated animals^6,7,16^, and suppress competing non-food-directed behaviors^10,17–19^. Recent studies also revealed that activity of AgRP neurons decreases rapidly upon sensory detection of food, before consumption begins, and shows anticipatory responses to food cues during innate and learned behaviors^8,9,20–23^. These observations have largely come from home-cage or cued-feeding paradigms in which food appears at fixed, easily accessible locations^6,7,21,22^. Foraging behavior, by contrast, is a spatial and dynamic problem: an animal must navigate a large environment, move between distant resources, approach a remembered food site, and decide when to leave^24–26^. Recently, ablation of AgRP neurons has been shown to impair the ability to associate spatial and contextual cues with food availability^27^. However, whether AgRP neuron activity tracks the spatial dimension of foraging and the animal’s position relative to a food source remains largely unexplored. If this were the case, it would expand the functional role of hypothalamic circuits, traditionally viewed as purely homeostatic, and suggest that motivational state and spatial context are already reflected at this early stage of foraging-related information processing. Determining whether AgRP neurons carry spatially relevant information is therefore critical for understanding how metabolic state interfaces with foraging and goal-directed navigation.

To address these questions, we used fiber photometry to record AgRP neuron activity while mice foraged freely in different spatial environments. We then combined activity mapping with a generalized linear model (GLM) to isolate the contribution of spatial distance to food from other behavioral and consummatory variables that shape AgRP neuron activity. Across environments, AgRP activity was predominantly explained by the animal’s distance to food after its discovery. This response was characterized by a dynamic, directional signal that fell on approach to the food location and rose on departure from it. Furthermore, we found that the AgRP distance signal was (i) regulated by the metabolic state, (ii) acquired through learning, and (iii) persisted during recall in a vision-dependent manner. Together, our findings suggest that AgRP neurons do more than report internal energy state or signal the sensory detection of food: their activity also reflects a learned spatial signal that links metabolic state to the memory of a food location. Our results broaden the scope of hypothalamic circuits beyond metabolic needs and consummatory behaviors, to reflect spatial information that could bias foraging toward remembered food sources.

## Results

### AgRP neuron activity during foraging signals spatial distance to food and metabolic state

To capture the activity dynamics of AgRP neurons during mouse foraging behavior, we expressed the calcium indicator GCaMP6s in AgRP neurons, using either Cre-dependent viral delivery (*AgRP^Cre^; AAV1-syn-FLEX-GCaMP6s*) or an intersectional transgenic strategy (*AgRP^Cre^>Ai162-TIGRE2.0-GCaMP6s*), and implanted an optical fiber over ARC for photometry recordings (Figures 1A and 1B, and S1A and S1C). Mice foraged in a maze with three arms (∼80 × 65 cm) in which the end of one arm contained an initially inaccessible compartment with food, allowing us to separate the period before food was available from the foraging epoch that followed door opening and food discovery (Figures 1C and 1D, Video S1). As expected, AgRP neuron activity was elevated in the fasted state and fell sharply at the moment of food discovery, remaining at a lower baseline across subsequent bouts of food visits and eating (Figures 1E, S1B and S1D). This drop was consistent across animals and GCaMP expression strategies, occurred at the onset of food discovery, and was accompanied by transient changes in AgRP activity dynamics during individual food visits (Figures S1E–S1H). Consistent with prior reports of state-dependent, anticipatory activity in these neurons^8,9,20,21^, suppression of AgRP activity at food discovery was greater in fasted mice than in *ad libitum*-fed mice (Figures S1E and S1F). Interestingly, we found that AgRP neuron activity was also dynamically regulated after food discovery, ramping down on approach and rising after food visits, in both fed and fasted animals (Figures 1E, S1G, and S1H).

**Figure 1.**
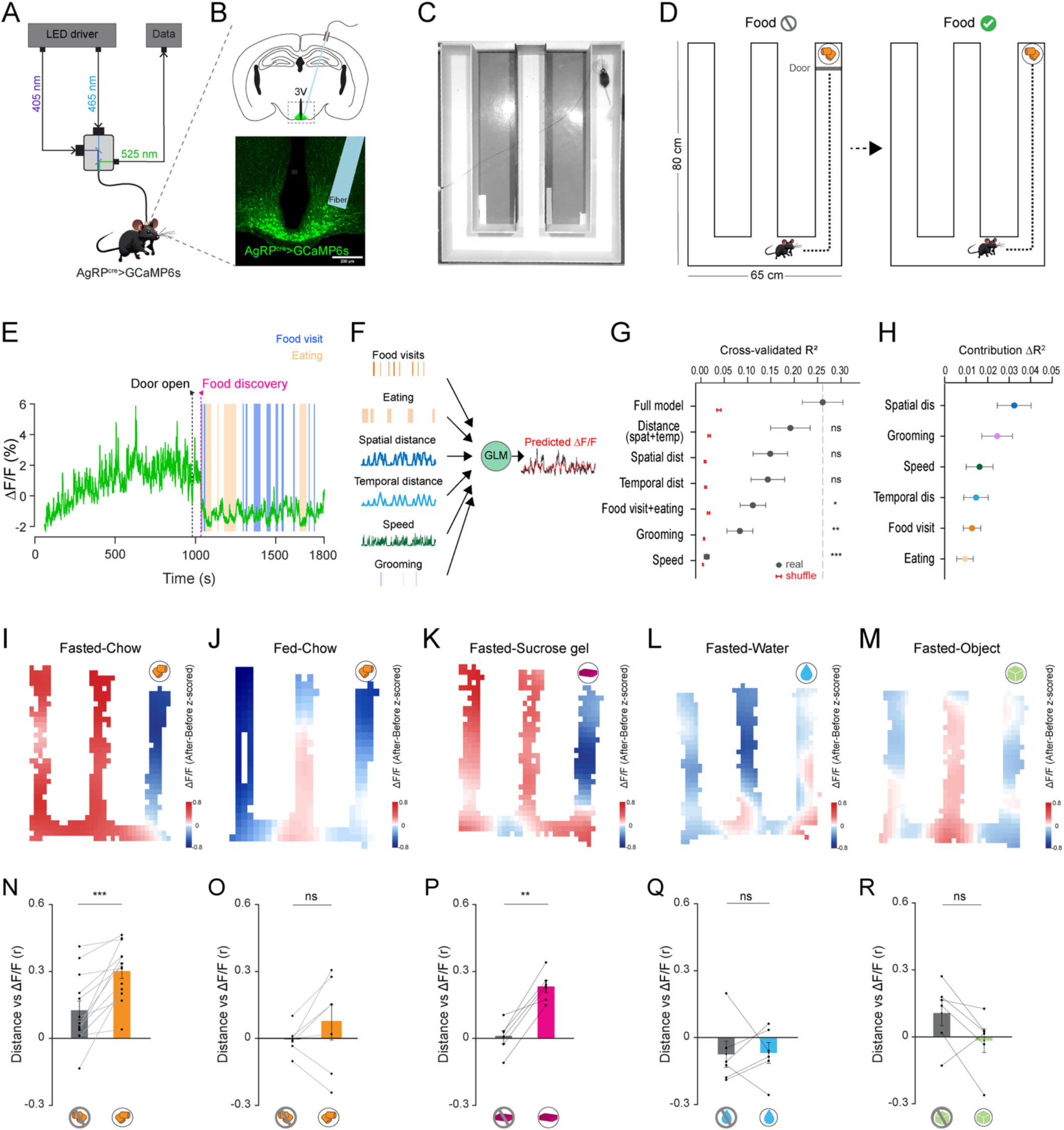
AgRP neuron activity correlates with spatial and temporal distance to a food source. (A) Fiber photometry configuration (465 nm excitation, 405 nm isosbestic, 525 nm emission). (B) GCaMP6s expression in AgRP neurons, with fiber placement (3V, third ventricle). (C) Top view of the three-arm maze showing the food location. (D) Foraging-task schematic. Note that the food compartment is separated from the maze by a solid door. Unavailable food = Food ⊘; available food = Food ✓. (E) Example AgRP neuron activity (ΔF/F) trace in a fasted mouse. Activity is high before access to food, drops at food discovery (magenta), and remains low across food visits (blue) and eating (orange). (F) Schematic of the GLM built for predicting AgRP neuron activity (ΔF/F) from multiple regressors, including food visits, eating, spatial/temporal distance, speed, and grooming. (G) Cross-validated R² for the full vs. reduced models (mean ± SEM; real vs. shuffled comparisons, One-way ANOVA followed by Dunnett’s multiple comparisons test; ns p>0.05, *p < 0.05; **p < 0.01; and ***p < 0.001). (H) Unique contribution to the explained variance (ΔR²) of each predictor (ns p>0.05, One-way ANOVA) (I–M) Maze heatmaps of activity change (after − before food discovery) during chow trials in fasted mice (I), chow trials in fed mice (J), sucrose gel trials in fasted mice (K), water trials in fasted mice (L), and object trials in fasted mice (M). (N–R) Correlation (r) between distance-to-food and AgRP neuron activity (ΔF/F) before and after food discovery in chow trials in fasted mice (N, n = 13 mice), chow trials in fed mice (O, n = 7 mice), gel trials in fasted mice (P, n = 6 mice), water trials in fasted mice (Q, n = 6 mice) and object trials in fasted mice (R, n = 6 mice). (**p < 0.01, ***p < 0.001; and ns, non-significant; paired t-test; Table S3 details statistical test and results).

To further dissociate among different behavioral variables that could contribute to the dynamic activity of AgRP neurons during foraging, we built a generalized linear model (GLM) to predict their activity (ΔF/F) from a set of regressors such as discrete food visits and eating bouts, locomotor speed, grooming, and the temporal and spatial distance to the food source (Figure 1F). The full model captured a substantial fraction of the AgRP variance (cross-validated R² above that of shuffled controls), and a reduced model containing only the distance terms (spatial + temporal) was statistically indistinguishable from the full model. On the contrary, models built from single behavioral predictors, including food visits, eating, grooming, or speed, performed significantly worse (Figures 1G, S2A–S2G). Spatial distance to food appeared to be the largest contributor among other unique regressors, although it was not statistically different from grooming, speed, temporal distance, food visits, or eating (Figure 1H). Because AgRP neuron activity can alter locomotor state^28^, we included speed as a predictor in the GLM, and the contribution of spatial distance persisted after accounting for it. Comparing the full GLM with models that excluded individual predictors confirmed that omitting the distance terms, particularly spatial distance, resulted in the largest decrease in predictive performance (Figure S2A-S2F, Table S2). Notably, this predictive structure emerged specifically after food became available (Figure S2H). These results suggest that, after food discovery, activity of AgRP neurons during foraging is better explained by the animal’s spatial distance to the food source than by food visits or consumption.

To visualize the spatial organization of this signal in detail, we mapped the change in AgRP neuron activity (z-scored) before vs. after food discovery across the three-arm maze. In 24-hr fasted mice foraging for standard chow, AgRP activity was strongly modulated by distance to the food, forming a clear gradient relative to the food location (Figures 1I and S3B). This gradient was absent when the same mice were fed *ad libitum* before the experiment (Figure 1J). To quantify this effect, we calculated the correlation between distance to food and AgRP activity for each animal. The distance–activity correlation increased significantly after food discovery in fasted mice foraging for chow, but not in fed mice (Figures 1N and 1O), indicating that this spatial modulation depends on metabolic state.

AgRP neurons are known to exhibit anticipatory activity and reflect food expectations based on sensory cues, including food odor^20^. Because odor in particular can diffuse across an environment and form spatial gradients, a distance-dependent AgRP signal could, in principle, arise from the animal sampling a food-odor gradient rather than from a learned representation of food location. We therefore asked whether sensory cues, and food odor specifically, were required for the distance-dependent AgRP activity to emerge, or whether it reflected the learned location of the food source. When fasted mice foraged for a palatable sucrose gel with minimal odor, AgRP neuron activity reflected the distance to the food, similar to what we observed with regular chow (Figures 1K, S3C, and S3D). Furthermore, the correlation between AgRP activity and distance to food increased significantly after gel discovery (Figure 1P), indicating that the AgRP activity gradient does not depend on strong olfactory cues and instead reflects the spatial proximity to the food. This distinguishes the signal from the rapid, consumption- or cue-driven decreases in other feeding paradigms and indicates that AgRP activity dynamics after food discovery are modulated by the animal’s distance from food rather than by its contact with the source.

We next investigated whether the AgRP distance-dependent activity reflects stimulus salience rather than the food reward itself. Both water and an object are behaviorally relevant to a foraging mouse. We presented fasted mice with water or an object in place of the food in the same foraging paradigm. In contrast to the chow and sucrose gel, the spatial activity maps did not show consistent proximity-related modulation for either stimulus, and the distance–activity correlation did not increase after water or object discovery (Figures 1L, 1M, 1Q, and 1R). Consistent with these results, the rapid suppression of AgRP activity seen for chow or sucrose-gel discovery was also absent at water and object discovery, and AgRP neuron activity was not regulated by distance from these stimuli (Figures S3E–S3H). Together, our results indicate that the distance-dependent modulation of AgRP activity is not driven by stimulus salience, but instead reflects the location of the available food source.

### AgRP neuron activity is dynamically regulated by approach and departure from food location

If AgRP neuron activity reflects distance to food during foraging, it should change gradually as the animal moves toward or away from the food source, rather than responding only at the moment of discovery or consumption. To test this prediction, we segmented foraging trajectories into “toward” (approach) and “away” (departure) runs, defined as relative distance to the food location (Figures 2A and 2B). Before food became accessible, runs toward or away from the chow or the sucrose-gel location were associated with changes in AgRP activity (Figures 2C, 2D, S3B, and S3D). After food discovery, however, AgRP neuron activity became strongly distance-dependent: during toward runs, activity ramped downward as the animal approached food, reaching a minimum near it, whereas during away runs, activity ramped back upward as the animal departed (Figure 2E, Video S1). The same bidirectional ramping was observed when fasted mice foraged for sucrose gel (Figure 2F), confirming that the effect generalizes across different food types. To quantify this relationship, we calculated the slope of AgRP activity (ΔF/F) versus distance to food for each run. Once chow or sucrose gel became available, approach (toward-run) slopes became significantly more negative and departure (away-run) slopes significantly more positive (Figures 2G–2J). This bidirectional, distance-dependent ramping activity indicates that AgRP neurons report not a static, all-or-none response to food but a continuously updated signal that tracks the animal’s instantaneous distance to the food, decreasing with proximity and increasing with separation. Notably, this graded activity cannot be explained by discrete consummatory or sensory-detection events alone, which would predict abrupt, step-like changes in activity locked to food contact rather than the smooth, direction-dependent ramps we observe. Instead, the same physical location was associated with opposite activity trajectories depending on whether the animal was approaching or leaving, indicating that AgRP activity reflects the animal’s location relative to the goal rather than its anticipation of the food.

**Figure 2.**
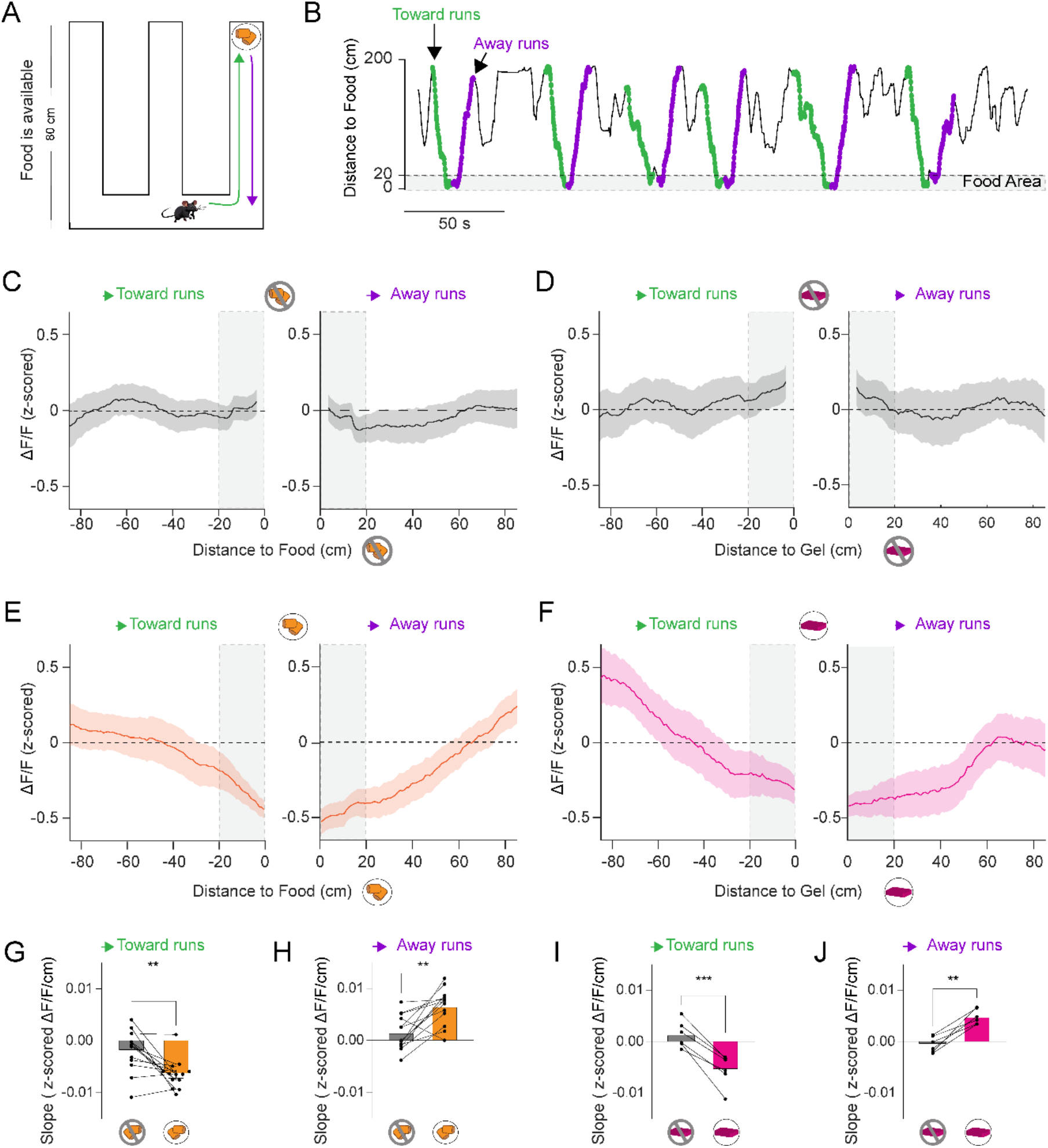
AgRP neuron activity decreases during food approach and increases during departure. (A) Schematic of toward (green) and away (purple) runs in the maze containing a single food source. (B) Example runs (distance to food vs. time); gray band indicates food area. (C–F) AgRP neuron activity vs. distance to food in indicated arms during toward (left) and away (right) runs before access to chow (C) and to gel (D), after access to chow (E), and to gel (F). (G–J) Slopes of the relationship between AgRP neuron activity and distance to food (ΔF/F/cm) when food was unavailable (before) vs. when it was available (after) for the chow trials and toward runs (G), chow trials and away runs (H), gel trials and toward runs (I) and gel trials and away runs (J) (n = 13 mice for the chow panels (G, H) and n = 6 mice for the gel panels (I, J); **p < 0.01, ***p < 0.001; paired t-test, Table S3 details statistical test and results).

### AgRP distance signal is acquired during learning and maintained during memory recall

Natural foraging relies on detecting available food sources and, in some cases, on remembering where a source was previously encountered. Having shown that AgRP neurons track proximity to food when a single source is present, we next asked whether accessibility and availability of food sources would interfere with this signal. For example, when several food sources share identical sensory properties, including food odor, but differ in accessibility, does AgRP neuron activity track the one source the animal can actually reach, or does it respond to all of them? These neurons also respond to learned cues previously associated with food^21^. Does the spatial gradient then persist selectively for a remembered food location once that food is no longer accessible? To address these questions, we recorded AgRP neuron activity in fasted mice while they foraged in a three-arm maze containing three food sources placed behind odor-permeable mesh doors at the end of each arm (Figure 3A). In this behavioral paradigm, mice were first allowed to freely explore the maze with no access to food, followed by a learning session during which one of the food sources became available at the end of one of the arms, and then tested in a final session two hours after learning without food access to assess memory recall. Mice initially explored all three arms equally but developed a robust spatial preference for the arm containing the accessible food during learning, which persisted during the memory recall test even though food was no longer accessible (Figures 3A–3C). We next quantified the dynamics of AgRP neuron activity across task phases. Spatial maps of AgRP activity during learning and memory recall, relative to the exploration phase, revealed a position-dependent reorganization in the maze arm containing the accessible food. The largest decrease in AgRP activity was localized to the available food source during learning (Figure 3D), and this reorganization was retained during the memory recall test (Figure 3E). Consistent with this, the distance-to-AgRP-activity correlation was indistinguishable among arms during the exploration phase, emerged strongly and selectively for the arm with the accessible food source during learning, and remained significant for that same arm during the memory recall test, when food was again inaccessible (Figures 3F–3H). We next quantified the approach and departure runs separately. During exploration, when all three odor-matched sources were present but inaccessible behind closed mesh doors, AgRP neuron activity showed no spatial modulation: toward- and away-runs to all three locations were flat, and approach and departure slopes did not differ across arms (Figures 3I, 3L, and 3O). This flat baseline suggests that the mere presence of food odor and the maze geometry are insufficient to generate the distance signal in AgRP neurons. Indeed, the activity gradient emerged only once the food source became accessible. During learning, AgRP neuron activity ramped downward as the animal approached the accessible food and upward as it departed, forming a spatial gradient that was absent at the two inaccessible food locations (Figures 3J and 3M), with significant approach and departure slopes only for the accessible arm (Figure 3P). Because all three sources emitted comparable food odor and differed only in accessibility, the selective emergence of the gradient for the single accessible source argues against olfactory cues, stimulus salience, or maze structure as sufficient explanations, and points instead to the learned location of the food source. Notably, this directional gradient persisted in the memory recall test. When all doors were closed again, and no food was accessible, AgRP neuron activity still ramped downward as mice approached the previously accessible location and upward as they departed, whereas the two never-accessible arms remained flat (Figures 3K and 3N). Approach and departure slopes for the remembered arm remained significant, though weaker than during learning, and exceeded those of the control arms (Figure 3Q). Thus, the bidirectional AgRP distance signal was not merely a response to food that was physically present; it could be recalled from memory and re-expressed during both approaches to and departures from the remembered food location.

**Figure 3.**
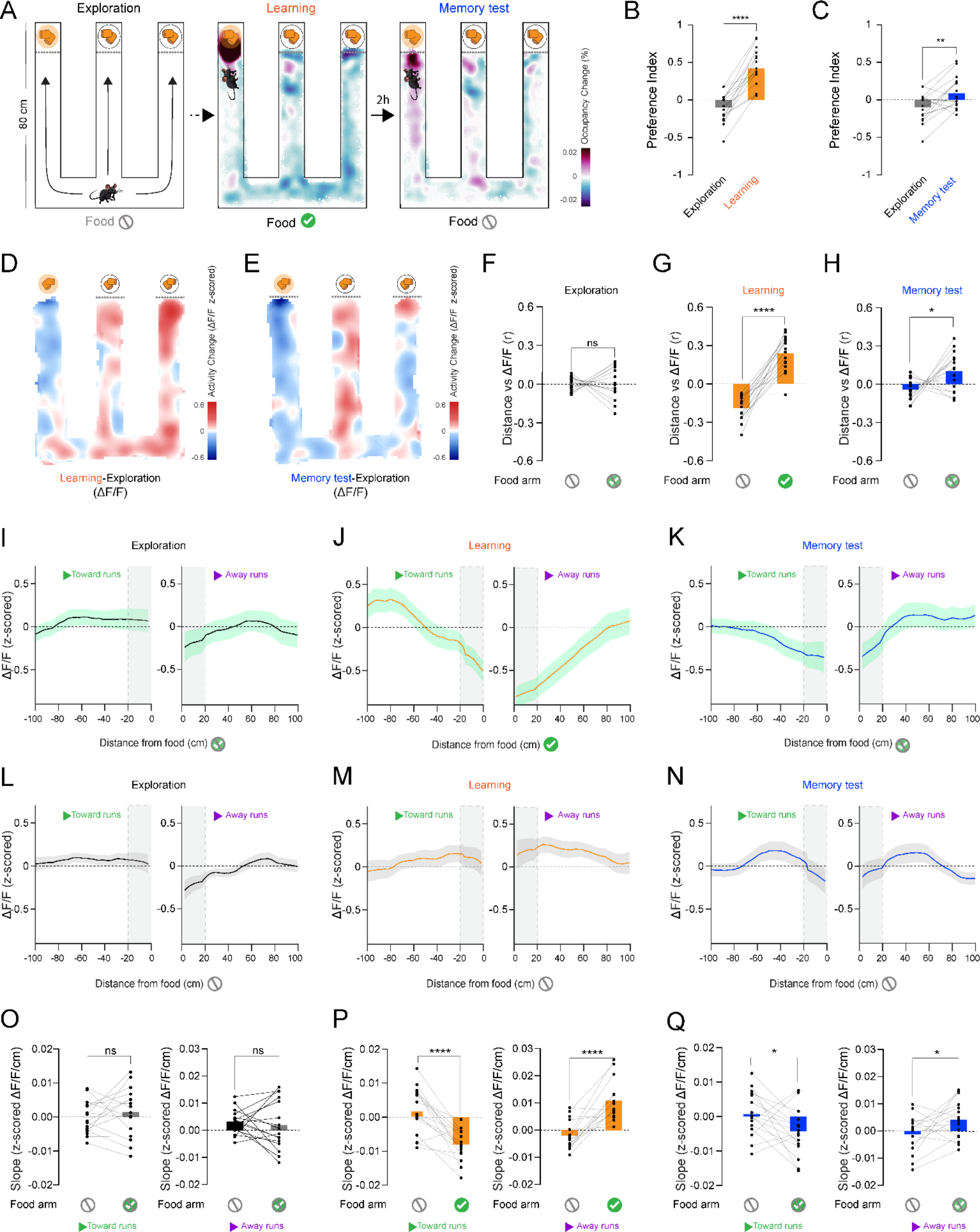
AgRP neuron activity reflects food availability and remembered food location. (A) Schematic of the three-arm maze paradigm with three food sources; one accessible during learning. Example occupancy-change heatmaps: exploration vs. learning (middle), exploration vs. memory recall test (right). (B) Preference index for the arm containing accessible chow during learning vs. exploration (****p < 0.0001, paired t-test, n=16). (C) Preference index for the previously accessible arm during the memory recall test vs. exploration (**p < 0.01, paired t-test, n=16). (D, E) Activity-change heatmaps (z-scored ΔF/F): learning − exploration (D), memory test – exploration (E) averaged across mice tested with the same available food arm (n=7). (F–H) Correlation (r) of AgRP neuron activity with distance to chow: control arms (no access to food) vs. food arm (accessible food) during exploration (F), during learning (G), during memory recall test (H) (n = 16 mice; *p < 0.05, ****p < 0.0001; ns; paired t-test). Arms: Food never available = ⊘; available food = ✓; food will be/was available =⊘+✓. (I–N) AgRP neuron activity toward (left) and away (right) runs vs. distance to food for the arm containing food (I–K) and the control arm (no food, L–N) across exploration (I, L), learning (J, M), memory recall test (K, N). (O–Q) Toward and away slopes in the control vs. food arm during exploration (O), learning (P), and memory recall test (Q) (n = 16 mice; *p < 0.05, ****p < 0.0001; ns; paired t-test).

Finally, we asked whether the AgRP distance signal depends on visual-spatial cues by repeating the paradigm under light and dark conditions (Figure S4). In these experiments, the spatial preference of mice and the AgRP neuron activity gradient developed during learning in both conditions, indicating that it does not strictly require vision (Figures S4A-S4N). Results during the memory recall test, however, differed between the two: In light conditions, both the behavioral preference and the distance modulation of AgRP neuron activity persisted during the memory recall phase, whereas in the dark, mice did not display a preference for the previously accessible food location, and AgRP activity was no longer correlated with the distance to it (Figures S4A-S4N). The parallel loss of both measures in the dark links the AgRP distance signal to spatial memory: when memory recall fails, the activity gradient is absent as well. Taken together, our results indicate that AgRP neuron activity reflects spatial proximity to the currently accessible food source, despite competing, equally odorous alternatives, and that distance-dependent AgRP activity is shaped by learning rather than solely by food odor.

### AgRP neuron activity reflects the distance to the food source rather than the maze geometry

To determine whether the AgRP learned distance signal requires the constrained geometry of a linear maze, we tested mice in a “cheeseboard” arena, a circular open field (∼76 cm diameter)^29^ containing 156 empty holes and no corridors in a similar foraging task (Figures 4A and 4B). Because this arena lacks the linear structure of the three-arm maze, AgRP neuron activity could, in principle, reflect space differently, or not at all. During an initial exploration phase, mice freely explored the arena with all holes empty. During the learning phase, a single hole was baited with a chow pellet, and mice learned its location over 10 trials, taking progressively shorter, more direct paths to the baited hole (Figures 4C and 4D). Similar to the three-arm maze (Figure 1), AgRP neuron activity decreased with proximity to the food, ramping down as the animal approached the baited hole during learning (Figures 4D and 4E). During the memory recall test (without food present), mice spent more time around the previously baited hole than during the initial exploration phase, and the correlation between AgRP activity and distance to the baited hole was elevated (Figures 4F–4I). We next analyzed the approach and departure runs separately. During memory-test approach runs, AgRP neuron activity decreased with proximity to the remembered target, forming a directional gradient that was absent in the exploration phase (Figures 4J and 4K). Departure runs, however, showed no significant correlation with distance (Figures 4L and 4M), unlike the three-arm maze, where both approach and departure runs were distance-tuned. This asymmetry is consistent with the difference between the two tasks: in the three-arm maze, the food remained physically present, though inaccessible, so its odor could continue to signal potential future availability in both directions, whereas in the cheeseboard, the baited hole was empty during the memory recall test, and mice had already discovered the absence of food, so the memory-driven signal was expressed predominantly during food approach but not departure.

**Figure 4.**
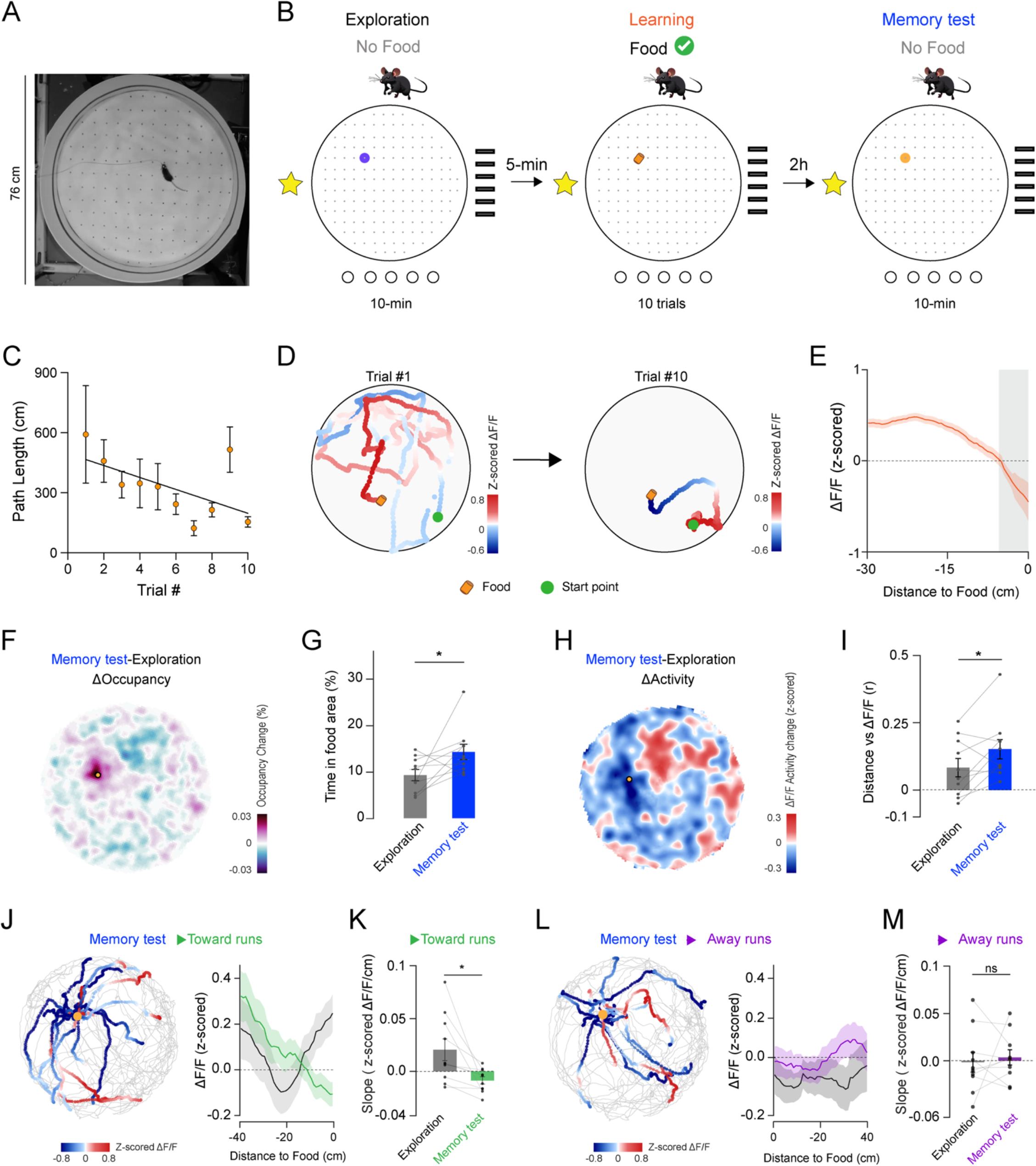
The AgRP food distance signal generalizes to an open arena with a hidden food source. (A) Top view of the cheeseboard maze (∼76 cm). (B) Behavioral paradigm: exploration (no food; purple circle marks the future food location), learning (food, 10 trials), memory recall test (no food, 2 h later; the orange circle marks the learned food location). (C) Path length decreases over learning (linear regression, n = 10 mice; R² = 0.056, p = 0.0177). (D) Example trajectories, trials #1 (left) and #10 (right), colored by z-scored ΔF/F. (E) AgRP neuron activity vs. distance to the food location during learning. Activity ramps down with proximity (n = 10 mice). (F) Occupancy difference, memory recall test − exploration (n = 10 mice). (G) Time in the food area during exploration (grey) vs. memory recall test (blue, n = 10; *p < 0.05, paired t-test). (H) Activity difference, memory test − exploration (n = 10 mice). (I) Correlation (r) of AgRP neuron activity with distance to the baited hole during exploration vs. memory recall test (n = 10 mice; *p < 0.05, paired t-test). (J) Toward (approach) runs during the memory recall test. Left, example trajectories colored by z-scored ΔF/F; right, AgRP neuron activity versus distance to food for exploration (black) and memory recall test (green) (shading indicates SEM). (K) Approach (toward-run) slope during exploration versus memory recall test; the slope becomes more negative during the memory recall test (n = 10 mice; *p < 0.05, paired t-test). (L) Away (departure) runs during the memory recall test. Left, example trajectories colored by z-scored ΔF/F; right, AgRP neuron activity versus distance to food for exploration (black) and memory recall test (purple) (shading indicates SEM). (M) Departure (away-run) slope during exploration versus memory recall test; the slope is unchanged (n = 10 mice; ns, paired t-test).

Together, our results demonstrate that modulation of AgRP neuron activity by distance to the food source is not restricted to linear mazes; it generalizes to a large, unstructured open arena and both currently learned and previously remembered food locations. These findings suggest that AgRP neurons do not merely signal the detection or anticipation of a food source, but also reflect a learned association with where nutrients were previously found. This association is retained, though attenuated, during memory recall, positioning these neurons as a candidate link between metabolic state and spatial navigation during foraging.

## Discussion

Here, we show that AgRP neurons signal the spatial distance to food during foraging. Using unbiased, model-based analysis, we found that spatial distance was the most significant correlate of AgRP neuron activity after food discovery, surpassing consumption and locomotion. The AgRP distance signal was directional and dynamic, ramping down on approach to the food and up on departure, and was regulated by metabolic state. Moreover, it was acquired through learning and retained in memory after food removal. These properties extend the role of AgRP neurons beyond signaling food anticipation and detection, suggesting that their activity reflects learned, experience-dependent information about the location of available food.

The AgRP spatial representation is most parsimoniously interpreted as a learned, value- and state-gated expectation signal rather than a primary spatial map. Previous work has shown that AgRP neurons display anticipatory activity^20,21^. Our results are consistent with this phenomenon but further reveal a spatial component to these responses during foraging. Whereas previously reported anticipatory decreases in activity occur on the order of a few seconds in response to discrete sensory cues, here the activity gradually decreased with proximity to food over tens of seconds during approach, whether or not the animal subsequently ate. Moreover, during departure, activity increased with distance from food, indicating that AgRP neurons track the spatial location of food rather than purely anticipate consumption. Notably, unlike previous paradigms where access to food was externally limited, here the mouse voluntarily disengaged from freely available food, and AgRP neuron activity increased as it moved away. This indicates that the increase in AgRP neuron activity accompanies self-initiated departure from freely available food, not only external changes in food availability. These results indicate that AgRP neurons provide a spatially modulated representation of food location that goes beyond momentary sensory cues or imminent consumption. A distance-related neuron signal of this kind has precedent in other systems: midbrain dopaminergic neural activity and striatal dopamine release ramp up as rodents navigate toward remote rewards, scaling with both the distance to and the value of the goal^30–32^. The AgRP signal reported here runs in the opposite direction, declining as the animal nears food, as expected of a hunger signal relieved by proximity to an anticipated meal rather than a reward-expectation signal that grows with it. A signal that decreases with proximity and increases with separation could bias the animal toward remembered food and influence the decision of when to leave, integrating physiological need with spatial expectation. Because this modulation is gated by metabolic state, it could provide a mechanism by which internal state can selectively recruit spatial memory of food, rather than of arbitrary locations. The selective expression of the distance signal in the open arena further indicates that AgRP dynamics are most strongly modulated during approach to the food source. Whether this distance-dependent signal is used to guide foraging behavior, and which circuits convey spatial information to AgRP neurons, remain to be determined. Two major inputs to AgRP neurons, the paraventricular hypothalamus (PVH) and the dorsomedial hypothalamus (DMH)^11,33^, are strong candidates. Recent work shows that the LH→DMH→AgRP pathway contributes to the anticipatory activity of AgRP neurons^22,34^. In parallel, prior studies have demonstrated that hippocampal spatial signals to the lateral septum (LS) are essential for foraging-related memory^35,36^. The LS projects to the lateral hypothalamus (LH)^37,38^, raising the possibility that a hippocampus→LS→LH→DMH→AgRP pathway could relay spatial information to AgRP neurons. Further work is needed to determine whether this circuit, or others, mediates the spatial modulation of AgRP neuron activity observed here and to identify the computations that occur along the pathway. Taken together, our data indicate that AgRP neurons in fasted mice reflect the spatial proximity of learned food locations. By showing that a homeostatic hunger circuit reflects and retains information about the distance to food, our findings position AgRP neurons as a candidate interface between internal state, spatial memory, and foraging behavior.

## Acknowledgements

We thank members of the Yapici, Oliva, and Fernandez-Ruiz laboratories, and Ruben Portugues for helpful discussions and critical feedback on the manuscript. We thank Marcelo de Oliveira Dietrich, Jesse Goldberg, and Ralitsa Todorova for helpful input on the experiments and analyses at the early stages of the project, and Renee Henderson for assistance with surgeries and behavior experiments. This work was supported by the National Institutes of Health (1R34NS128872 to N.Y., R01MH140009 to A.O), a Cornell Neurotech Mong Fellowship (to A.G.), and a Sigma Xi Cornell Fellowship (to A.G.).

## Author contributions

A.G. and N.Y. conceived the study and designed the overall research framework. A.G., N.Y., A.O., and A.F.R. designed the behavioral and recording experiments. A.G. and J.S. performed stereotaxic surgeries, fiber photometry recordings, and behavioral experiments, and collected the fiber photometry and position-tracking data. A.G. developed and implemented the analysis pipeline, including generalized linear modeling, distance-encoding, slope analyses, and spatial activity maps. D.S. contributed to data preprocessing, signal extraction, and behavioral tracking. A.G. led the interpretation of the results and, together with N.Y., wrote the manuscript. All authors discussed the results, provided critical feedback, and approved the final version of the manuscript.

## Declaration of Generative AI Use

During the preparation of this work, the authors used Claude (Anthropic) and ChatGPT (OpenAI) to edit code and improve the language, clarity, and readability of the manuscript. After using these tools, the authors reviewed and edited the content as needed and took full responsibility for the content.

## STAR ★ Methods

### Resource Availability

#### Lead contact

Further information and requests for resources and reagents should be directed to and will be fulfilled by the lead contact, Nilay Yapici.

#### Materials availability

This study did not generate new, unique reagents.

#### Data and Code availability

- Raw fiber photometry and mouse behavioral data are deposited on servers at Cornell University and are available upon request by contacting the lead contact.
- Animal genotypes, statistical tests, and data in the figures are available as supplementary tables.
- The original code that supports the findings of this study can be downloaded from GitHub (https://github.com/Nilayyapici/Gruzdeva-et-al).

### Experimental model and subject details

Mice used in this study included *AgRP-cre* mice: *Agrp^tm1(cre)Lowl^/J* (Jackson Laboratory, stock no. 012899), and *AgRP-cre>GCaMP6s* mice were generated by crossing *B6.Cg-Igs7^tm1<u>62</u>.1(tetO-GCaMP6s,CAG-tTA2)Hze^/J* (stock no. 031562) with *Agrp^tm1(cre)Lowl^/J*. Both male and female mice, aged 2–10 months, were housed in groups of 1–5 before and after surgery under a 12-hour reverse light/dark cycle. All experiments conformed to guidelines established by the National Institutes of Health and have been approved by the Cornell Institutional Animal Care and Use Committee. Detailed genotypes are listed in Table S1. The study was not powered to detect sex differences, and data were pooled from both sexes.

## Method details

### Surgery and viral injections

Mice were anesthetized with isoflurane and administered buprenorphine (0.05 mg/kg, subcutaneously) and dexamethasone (5 mg/kg, subcutaneously) for analgesia and control of inflammation. Animals were head-fixed in a stereotaxic apparatus (David Kopf Instruments) and injected with lidocaine (0.2%) for local analgesia. *AgRP-cre* mice received bilateral injections of *AAV1.Syn.Flex.GCaMP6s.WPRE.SV40* (2.1 × 10^13 GC/mL) or *AAV9.hSyn.DIO.jGCaMP8s.P2A.ChrimsonR.ST* (2.5 × 10^13 GC/mL) (mice injected with this construct were used in Figures S4A–S4N) into the ARC. Two 1-mm diameter craniotomies were made at coordinates AP: –1.65 mm, ML: ±0.25 mm relative to Bregma. A glass pipette (3”, Drummond Scientific) backfilled with mineral oil and mounted on a Nanoject III (Drummond Scientific) was used to deliver the virus (four injections (150 nl each): AP: –1.55 mm, ML: ±0.25 mm, DV: 5.75 mm; and AP: –1.65 mm, ML: ±0.25 mm, DV: 5.75 mm). One to two weeks after viral injection, mice were implanted with an optical fiber (R-FOC-L200C-39NA, 6 mm length, RWD) targeting the ARC (coordinates: AP: –1.6 mm, ML: 0.25 mm, DV: 5.72 mm). *AgRP-cre>GCaMP6s* mice did not receive viral injections but were implanted with the optical fiber using the same procedure. After each surgery and for the following two days, mice received ketoprofen (5 mg/kg, subcutaneously) for post-operative analgesia. Fiber photometry recordings were conducted in freely moving mice after a recovery period of at least three weeks. After the experiments, mice were perfused, and GCaMP6s expression in the AgRP neurons was confirmed on brain slices. Virus injections per mouse are listed in Table S1.

### Fiber photometry

Fiber photometry recordings were performed using a Doric Lenses system consisting of a Fiber Photometry Console (FPC), LED driver, and integrated Fluorescence Mini Cube (ilFMC4), controlled via Doric Neuroscience Studio software. Excitation light at 405 nm (isosbestic control) and 465 nm (GCaMP excitation) was delivered through a patch cord with a rotary joint, with light power at the fiber tip adjusted to 60 μW for 465 nm and 14 μW for 405 nm. Signals were acquired in lock-in mode, with modulation frequencies of 208.6 Hz for the 405 nm LED and 572.2 Hz for the 465 nm LED.

### Fasting

Animals were tested either ad libitum or after a 24-hour fast. For fasting, all food was removed from the home cage for 24 hours, while access to water remained ad libitum. Following a fasting session, animals were not fasted again until at least one full day had passed. The start time of fasting varied between 11:00 a.m. and 4:00 p.m., depending on the timing of the experiments.

### Experimental environment and video acquisition

Experiments were conducted under dim white light, except for the dark-condition sessions (Figures S4D–S4F and S4K–S4N), in which visible light was extinguished, and behavior was recorded under infrared illumination only. To support spatial orientation, high-contrast visual landmarks were placed on the walls and floor of the three-arm maze and at fixed locations in the surrounding environment, providing both local and distal spatial cues. Behavior was recorded with two Logitech webcams (C920 HD Pro and C922 HD Pro) at 1024 × 576 resolution and a 30 Hz frame rate: one camera was mounted above the mazes to track the animal’s position (top view), and a second camera was positioned over the food dish to monitor feeding behavior in the three-arm maze (food camera). Two infrared (IR) light sources (850 nm) above the mazes provided illumination for video tracking, including in darkness; to improve IR sensitivity, the IR-blocking filters were removed from the webcams. AgRP neuron population activity was recorded using fiber photometry, and photometry streams (60 Hz) were acquired and synchronized with Bonsai (version 2.8.2) and an Arduino microcontroller. Photometry (60 Hz) and video tracking (30 Hz) were synchronized by interpolating behavioral variables (position, speed) to photometry timestamps using linear interpolation, then binned into 0.2 s bins (one source maze) or 0.05 s bins (three sources maze and cheeseboard) by averaging photometry signals and taking the end-of-bin value for positional variables.

### Behavioral tests

#### Three-arm maze with one source

Mice were allowed to freely explore a modified three-arm maze^39^ (external dimensions: total width: 65cm; arm length: 80 cm; arm width: 10 cm; wall height: 12 cm) containing a single food source placed in a plastic dish at the end of one of the arms (the specific arm varied across mice). For the first 15 minutes, the food source was kept behind a solid plastic door to prevent access. In the following 15 minutes, the door was manually removed, allowing free access to the food. Two food sources were used: a standard home-cage chow pellet soaked in water to soften it (dry weight=2.5 g), and an odorless hydrogel containing 1M sucrose (3 g of the gel). For non-food (control) trials, a water bottle (height 6.9 cm, base: 8 cm in diameter, bottle (top) 3.8 cm in diameter) or an object (wooden cylinder, height 6.9 cm, diameter 4.8 cm) was presented at the food location in place of the food. Water and the object served as non-food controls to test the specificity of AgRP neuron responses to food (Figures 1Q, 1R, and S3E-S3H). To minimize neophobia, both foods, the water bottle, and the object were introduced to the mice in their home cages before the start of experiments, so that the stimuli were familiar at the time of testing.

#### Three-arm maze with multiple food sources

24-hour-fasted mice explored a three-arm maze with multiple food sources; each was placed at the end of a different arm. All food sources were soaked pellets from the home cage. During the first 10 minutes (exploration), the food dishes were placed behind mesh-covered doors to allow food odors to diffuse without access. During learning, the door to one arm (varied across mice) was opened for 4 minutes and closed for 2 minutes, repeated four times, while two other arms contained inaccessible food sources (control arms). After the learning session, mice were returned to their home cages without access to food. Two hours later, mice were tested for 10 minutes in the same three-arm maze with all mesh doors closed (memory test).

#### Cheeseboard maze

In the initial session (exploration), mice were placed in the empty circular cheeseboard maze (76 cm diameter) containing 156 evenly spaced holes (5 mm in diameter)^29^ on the south side and allowed to explore freely for 10 minutes. Five minutes after the exploration session, mice underwent a learning session. During the learning session, a single hole (varied across mice) was baited with a 0.03–0.06 g piece of food pellet from the home cage. For each of the 10 trials, mice were placed into the arena from one of four directions (south, east, north, or west) in a pseudorandom order. Once a mouse located and consumed the food, it was removed from the arena for 1–2 minutes before the next trial. If the mouse did not find the food within 4 minutes, the trial was terminated. Experiments were stopped if a mouse failed to complete more than three consecutive trials. After the learning session, mice were returned to their home cages. Two hours after the learning session, mice were tested in a 10-minute probe trial in the same, now-empty cheeseboard maze (Memory test).

### Tissue processing and histology

After completion of the experiments, mice were deeply anesthetized with sodium pentobarbital (100 mg/kg, Vortech Pharmaceuticals, #VPL9373) and perfused with 1x phosphate-buffered saline (PBS), followed by 4% paraformaldehyde (PFA) in PBS. Brains were extracted and post-fixed in 4% PFA for 24 hours at 4 °C, then cryoprotected in 30% sucrose in PBS at 4 °C until they sank (≥24–48 hours). Coronal sections (50 µm) were cut on a sliding microtome (Leica SM2000R) and collected as free-floating sections in PBS. Sections containing the ARC were mounted onto Superfrost Plus slides and coverslipped with mounting medium (VECTASHIELD with DAPI to counterstain nuclei). Sections were imaged on a fluorescence microscope (Thorlabs Cerna with a Zeiss 10× objective) to verify GCaMP expression and optic-fiber placement.

## Quantification and statistical analysis

### Fiber photometry

To calculate ΔF/F from the fiber photometry signal, a least-squares linear fit was first applied to the 405 nm signal to align it with the 465 nm signal. The fitted 405 nm trace was then used to normalize the 465 nm signal according to the following formula: ΔF/F = (465 nm signal − fitted 405 nm signal)/fitted 405 nm signal^40^. To quantify directional changes in neural activity relative to food location, we implemented a spatial slope analysis on individual movement runs. Runs were defined as directed movements toward or away from food sources. Distance from a source was computed from tracked x/y coordinates and smoothed with a first-order low-pass Butterworth filter. The temporal derivative of each distance signal was used to classify movement direction. A towards run began when the distance derivative fell below −0.1 while the animal was within the food area threshold (20 cm from food) and was retained only if the animal reached the arm endpoint. An away run began when the derivative exceeded +0.1 from within the food area and was retained only if the animal had been near the arm at run onset. Both run types terminated when the derivative crossed back through its respective threshold. For each mouse, ΔF/F signals were first z-scored within session to normalize for individual baseline differences. For the cheeseboard maze, the same derivative-based algorithm was applied to the distance from a single food-hole location, with a food-area threshold of 6 cm and a minimum run duration of 0.5 s. Group-level inference was conducted using a mixed-effects approach: a separate linear regression was first fit for each mouse, yielding one slope estimate per mouse per condition.

### Generalized linear model

A generalized linear model (GLM) predicted z-scored ΔF/F from food visits, eating, locomotor speed, grooming, temporal distance (time since food discovery), and spatial distance to food (Figure 1F). Prior to fitting, ΔF/F and all continuous predictors (speed, spatial distance, and temporal distance) were z-scored. Models were evaluated using cross-validated R² (CV-R²) (Figure 1G) and mean squared error (MSE) (Figure S2G). Cross-validation used 100 iterations of random 80/20 train/test splits, with cross-validated R² (CV-R²) computed on held-out frames and averaged across iterations. Shuffle controls applied circular shifts to the ΔF/F trace by a random offset (10–90% of the session length), preserving the autocorrelation structure of the signal while destroying its alignment with behavioral predictors; shuffle R² was averaged over 20 independent shifts per session. Fifteen models were compared: a full model containing all predictors, leave-one-out ablations (removing spatial distance, temporal distance, both distance measures, food visits, eating, food visits and eating combined, grooming, or speed), and single-predictor isolation models (spatial distance only, temporal distance only, both distance measures, food events only, grooming only, and speed only) (Table S2). The unique variance attributable to each predictor (ΔR²) was computed as the reduction in adjusted R² when that predictor was removed from the full model (Figures 1G, 1H, and S2A-S2G). Group-level significance was assessed using a one-way ANOVA, and comparisons with the full model were assessed with Dunnett’s multiple-comparison tests. Before/after comparisons were made using a paired t-test, comparing the pre-food-availability session (before) and the post-discovery session (after) to assess whether predictive structure emerged upon food availability.

### Behavioral quantification

The mouse’s nose position was tracked from the overhead video using DeepLabCut^41^ (version 2.3.5). Tracked coordinates were used to compute the animal’s instantaneous position, locomotor speed, and spatial or temporal distance to the food location. Eating and non-eating food visits were manually annotated from close-up video of the food dish, and grooming episodes were manually scored from overhead video of the entire maze by trained observers, blind to experimental condition. Grooming periods were excluded from the distance-based analyses because AgRP neuron activity increased during grooming; however, grooming was retained as a nuisance predictor in the GLM. All subsequent analyses were performed using custom MATLAB code (MathWorks, version R2024a). AI-assisted tools (ChatGPT, OpenAI; Claude, Anthropic) were used to help draft and debug this analysis code; the authors reviewed, tested, and validated all code.

#### Preference index

Spatial preference for the food-associated arm was quantified from positional occupancy in each session (exploration, learning, and memory recall test). The food arm was defined as the arm containing accessible chow during learning; the same physical arm was used across all three sessions, and all arms were inaccessible during exploration and the memory recall test. For each session, we computed *T*_food_, the time the animal’s tracked position fell within the food arm, and *T*_ctrl_, the mean occupancy time across the remaining (never-baited) arms. The preference index was then calculated as

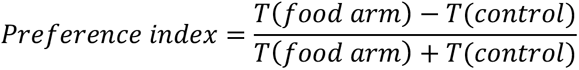

This index ranges from −1 to +1, where +1 indicates exclusive occupancy of the food arm, 0 indicates no preference among arms, and negative values indicate avoidance of the food arm. Preference indices were compared between sessions (exploration vs. learning; exploration vs. memory recall test) using two-tailed paired t-tests (Figures 3B, 3C; Figures S4B, S4C, S4E, and S4F).

### Statistical tests

Statistical analyses were performed in GraphPad Prism (version 11.0.2). Comparisons between two paired groups were made using two-tailed paired t-tests. Comparisons among more than two groups were made using ordinary one-way ANOVA followed by Dunnett’s multiple comparisons test. The relationship between path length and trial number was assessed by simple linear regression. All tests were two-tailed, and statistical significance was defined as p < 0.05. Significance levels are indicated as follows: *p < 0.05, **p < 0.01, ***p < 0.001, and ****p < 0.0001. Data are presented as mean ± SEM unless otherwise indicated. The statistical test, exact p-value, and sample size (n) for each comparison are reported in the corresponding figure legend and in Table S3.

**Video S1. AgRP neuron activity during foraging in a three-arm maze with a single food source, related to Figure 1**. Recording from a 24-hour-fasted mouse foraging for soaked chow. Left, top-view camera; right, food-zone camera. AgRP neuron activity (% ΔF/F, green) and distance to food (cm, magenta) are plotted below. Food interactions, eating, and grooming bouts are labeled in the bottom-left of the top-view panel. The video is sped up 2×.

## Supplementary Figures, Figure Titles, and Legends

**Figure S1.**
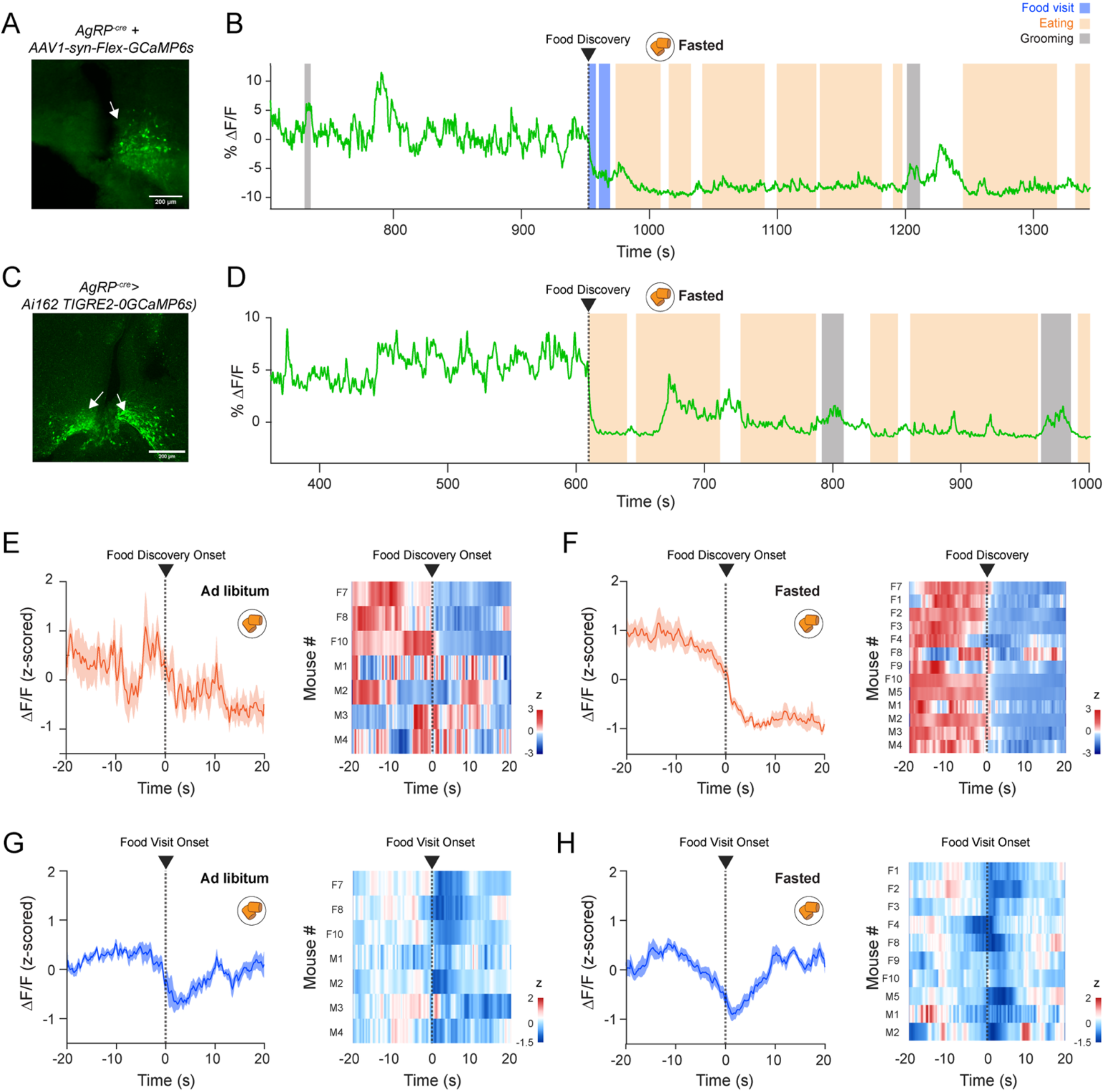
AgRP neuron activity decreases at food discovery and food visits in fasted and ad libitum mice (Related to Figure 1). (A, C) Histological verification of GCaMP6s expression in the arcuate nucleus (ARC) for *AgRP-Cre* mice injected with *AAV1-syn-FLEX-GCaMP6s* (A) and *AgRP-Cre>Ai162-TIGRE2.0-GCaMP6s* mice (C). (B, D) Example photometry recordings with annotated behaviors (blue, food visits; orange, eating; gray, grooming) for the two strategies described in A and C. (E, F) AgRP activity (ΔF/F) aligned to food-discovery onset (left) with per-mouse average heatmaps (right): ad libitum (E, n = 7 mice) and fasted (F, n = 13 mice). (G, H) AgRP activity (ΔF/F) aligned to food-visit onset (left) with per-mouse average heatmaps (right) in *ad libitum* (G, n = 7 mice) and fasted (H, n = 10 mice).

**Figure S2.**
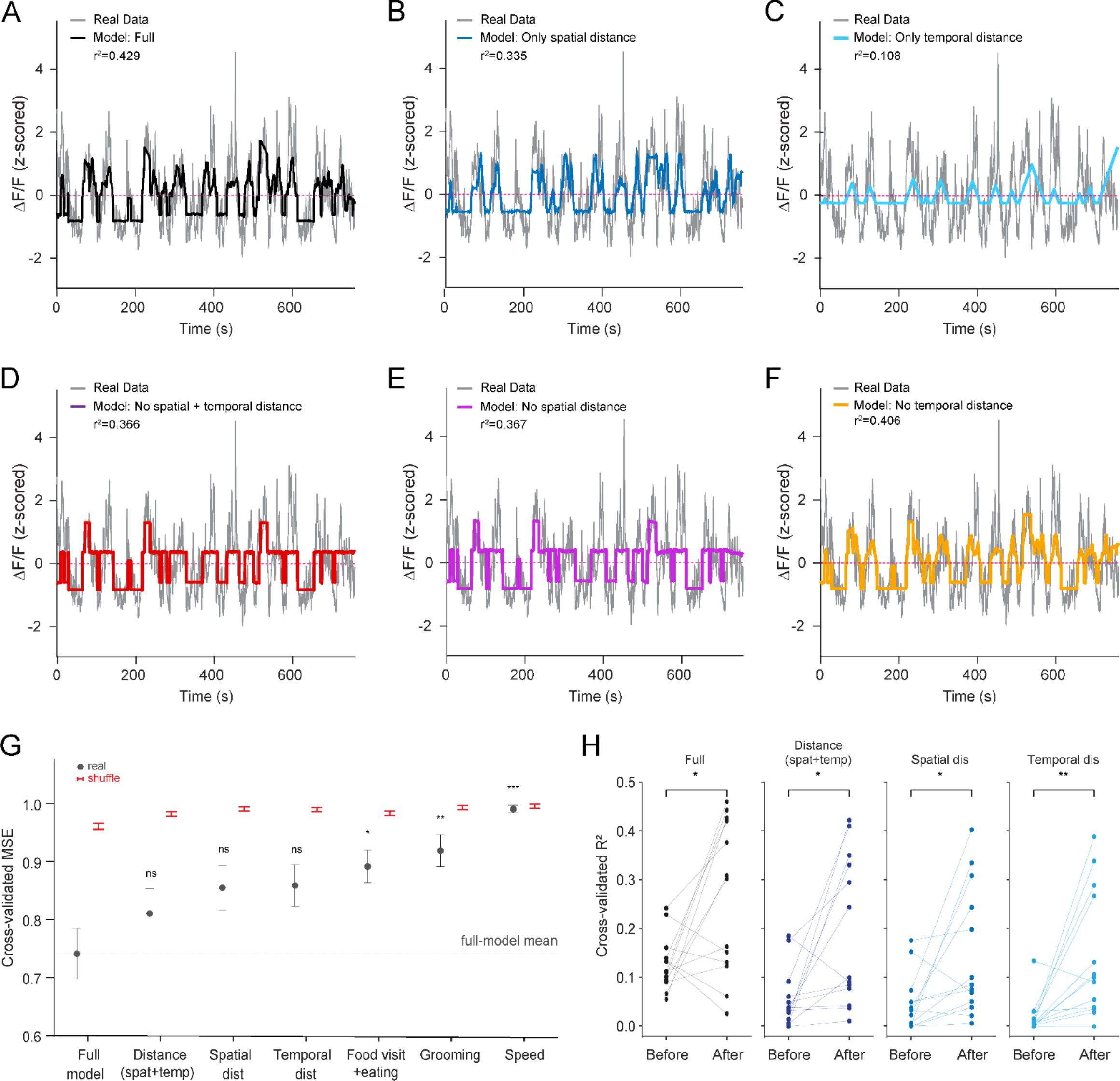
Spatial distance is a significant predictor of AgRP neuron activity in the generalized linear model (Related to Figure 1). (A–F) Example predicted (colored) vs. observed (gray) AgRP activity (ΔF/F) for different GLMs: full model (A), spatial distance only (B), temporal distance only (C), excluding spatial and temporal distance (D), excluding spatial distance (E), excluding temporal distance (F). (G) Cross-validated mean squared error (MSE) per model vs. shuffled controls (predictors circularly shifted). Model comparisons relative to the full GLM (p> 0.05, ns.; *p < 0.05, **p < 0.01, and ***p < 0.001 for food visit and eating, grooming, and speed, respectively; one-way ANOVA followed by Dunnett’s multiple comparisons test; Table S3 details statistical test and results). (H) Cross-validated R² before vs. after food discovery for the full GLM, distance (spatial + temporal), spatial-only, and temporal-only models; performance increased after discovery (*p < 0.05, **p < 0.01; paired t-test).

**Figure S3.**
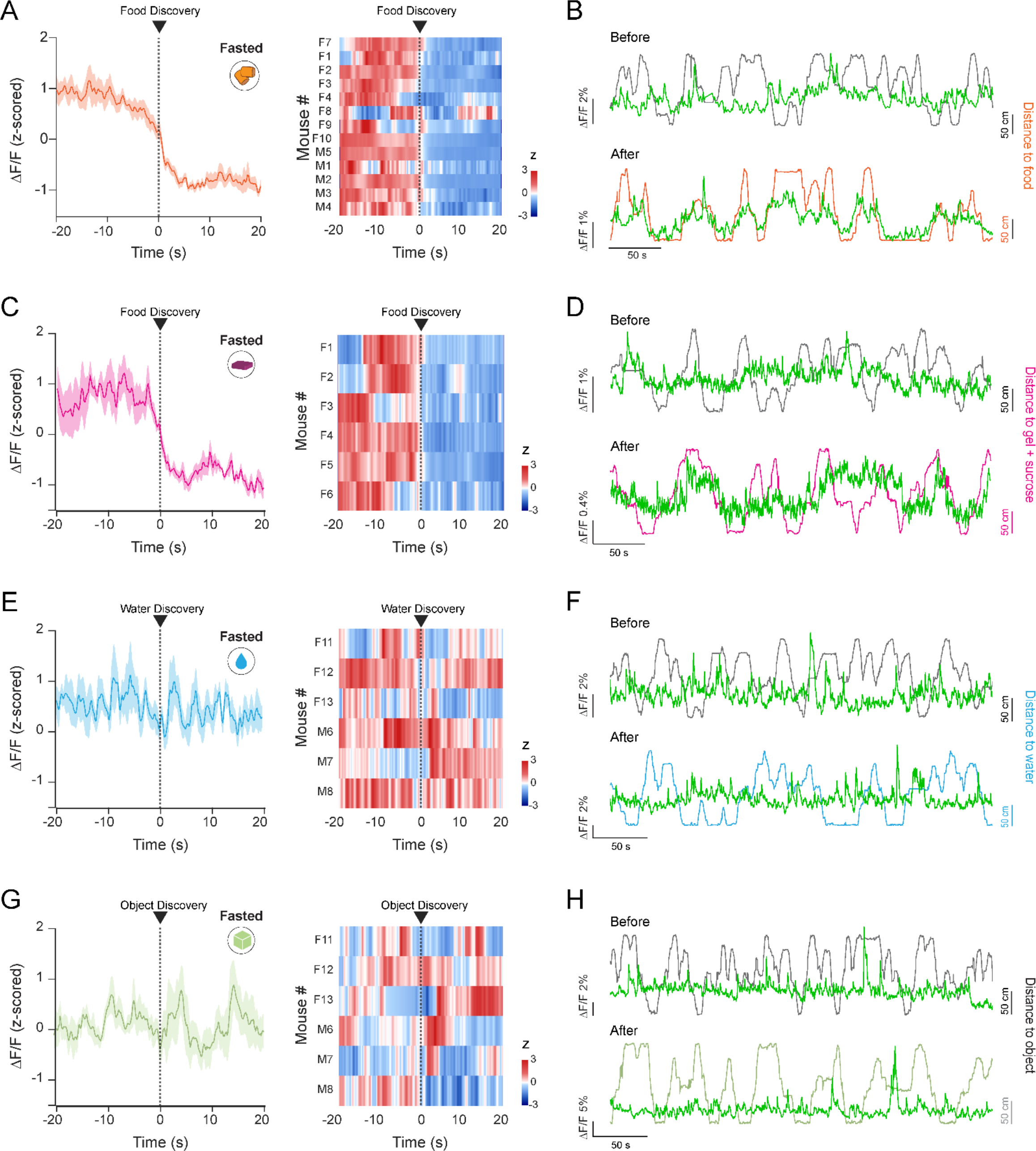
AgRP neuron activity tracks the discovery of and distance to food, but not water or an object (Related to Figure 1). (A, C, E, G) AgRP activity (ΔF/F) aligned to food discovery onset (left) with per-mouse average heatmaps (right) in fasted mice for chow trials (A, n = 13 mice, same as Figure S1F), sucrose gel trials (C, n = 6 mice), water trials (E, n = 6 mice), and object trials (G, n = 6 mice). (B, D, F, H) Example AgRP activity traces vs. distance to chow (B), sucrose gel (D), water (F), and object (H), before (top) and after (bottom) item discovery.

**Figure S4.**
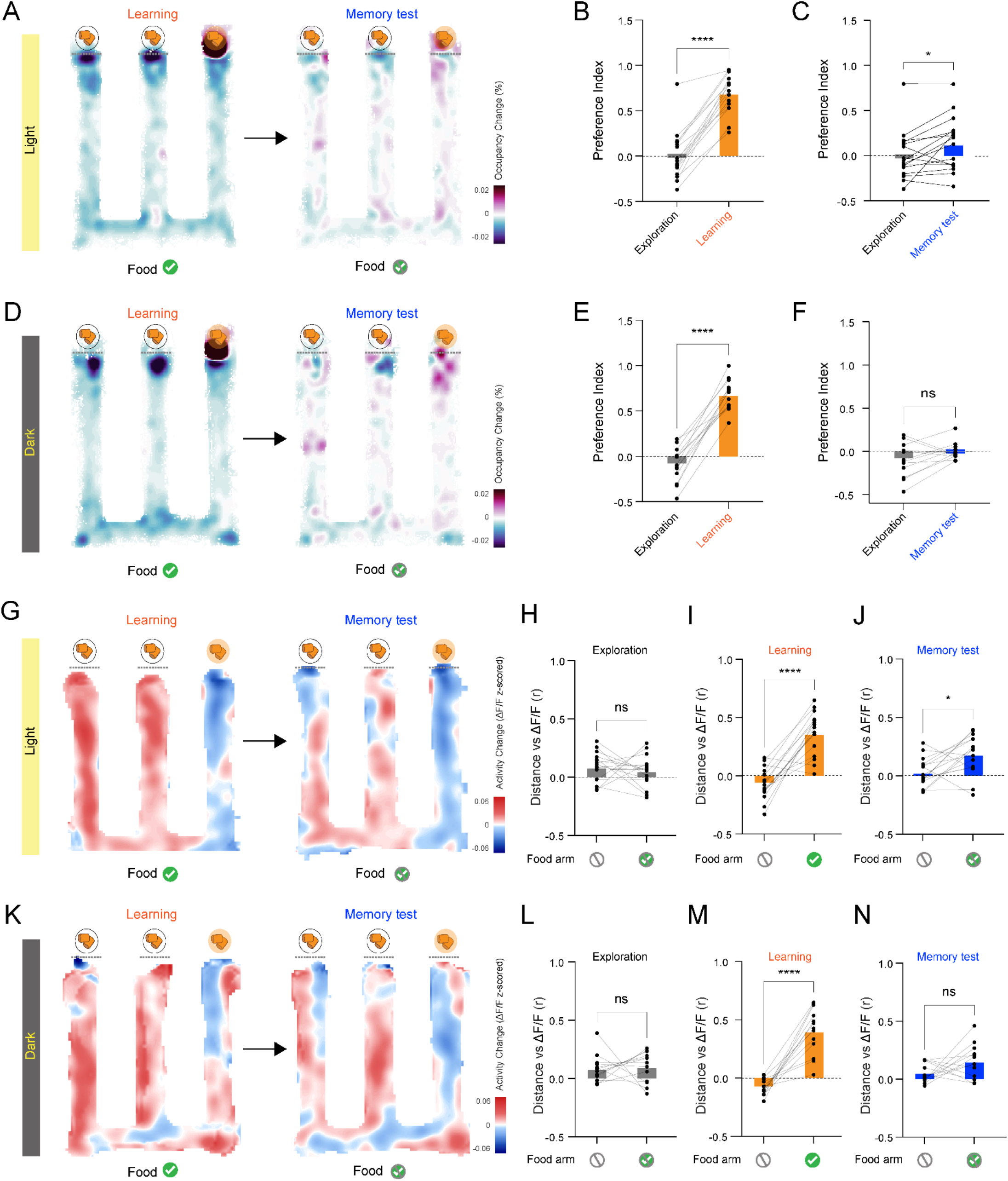
AgRP distance signal is acquired in both light and darkness but retained only in light during the memory test (Related to Figure 3). (A, D) Occupancy-change heatmaps during exploration vs. learning trials (left) and exploration vs. memory recall test (right), for mice with accessible chow in the right arm during learning, under light (A) and dark (D) conditions. (B) Preference index for learning vs. exploration during light sessions (n = 16 sessions from 8 mice; ****p < 0.0001; paired t-test). (C) Preference index for memory recall test vs. exploration during light sessions (n = 16 sessions from 8 mice; *p < 0.05; paired t-test). (E) Preference index for learning vs. exploration during dark sessions (n = 13 sessions from 7 mice; ****p < 0.0001; paired t-test). (F) Preference index for memory recall test vs. exploration during dark sessions (n = 13 sessions from 7 mice; ns; paired t-test). (G, K) AgRP activity-change heatmaps: learning − exploration (left) and memory recall test − exploration (right), for right-arm trials under light (G) and dark (K) sessions. (H–J) Correlation (r) of AgRP activity with distance to chow during exploration (H), learning (I), and memory recall tests (J), in light sessions (n = 16 sessions from 8 mice; ns, *p < 0.05 and ****p < 0.0001 for learning and memory recall test comparisons; paired t-test. Unavailable food = Food ⊘; available food = Food ✓). (L–N) Correlation (r) of AgRP activity with distance to chow in the arm containing the food during exploration (L), learning (M), and memory recall test (N) in dark sessions (n = 13 sessions from 7 mice; ns for exploration (L) and memory recall test (N), ****p < 0.0001 for learning (M); paired t-test).

